# Free Amino Acids Accelerate the Time-Dependent Inactivation of Rat Liver Nucleotide Pyrophosphatase / Phosphodiesterase Enpp3 elicited by EDTA

**DOI:** 10.1101/2024.06.27.601032

**Authors:** Ana Romero, Guadalupe Cumplido-Laso, Ascensión Fernández, Javier Moreno, José Canales, Rui Ferreira, Juan López-Gómez, João Meireles Ribeiro, María Jesús Costas, José Carlos Cameselle

## Abstract

Nucleotide-pyrophosphatases/phosphodiesterases (NPP/PDE) are membrane or secreted Zn^2+^-metallohydrolases of nucleoside-5’-monophosphate derivatives. They hydrolyze, for instance, ATP and 4-nitrophenyl-dTMP, and belong to the ecto-nucleotide pyrophosphatase/phosphodiesterase (ENPP) family that contains seven members (ENPP1-ENPP7). Earlier we had shown that an NPP/PDE activity solubilized and partially purified from rat liver membranes is inactivated by EDTA in a time-dependent fashion, an effect enhanced by glycine and blocked by the 4-nitrophenyl-dTMP. Here, we extended this observation to other free amino acids. Activity assays started after different incubation lengths with EDTA provided first-order, apparent inactivation constants (k_i(ap)_). With the exception of cysteine (a strong inhibitor) and histidine (itself evoking a time-dependent inactivation), free amino acids themselves did not affect activity but increased k_i(ap)_. The results are compatible with a conformational change of NPP/PDE evoked by interaction with free amino acids. The enzyme preparation was analyzed to identify what ENPP family members were present. First, the hydrolytic activity on 2’,3’-cGAMP was assayed because until very recently ENPP1 was the only mammalian enzyme known to display it. 2’,3’-cGAMP hydrolase activity was clearly detected, but mass spectrometry data obtained by LC-MS/MS gave evidence that only rat Enpp3, Enpp4 and Enpp5 were present with low abundance. This finding coincided in time with a recent publication claiming that mouse Enpp3 hydrolyzes 2’,3’-cGAMP, and that Enpp1 and Enpp3 account for all the 2’,3’-cGAMP hydrolase activity in mice. So, our results are confirmatory of Enpp3 activity towards 2’,3’-cGAMP. Finally, the effect of amino acids could be relevant to NPP/PDE actions dependent on protein-protein interactions, like the known insulin-related effects of ENPP1 and possibly ENPP3.

## Introduction

Nucleotide pyrophosphatases/phosphodiesterases (NPP/PDE) are membrane or secreted metalloenzymes that hydrolyze (near) all sort of phosphoanhydride and phosphodiester derivatives of nucleoside 5’-monophosphates (NMP), yielding NMP products (Borza et al. 2021; Stefan et al. 2005). The NPP/PDE preparation studied here was solubilized and partially purified from rat liver membranes (RLNPP/PDE), and is known to be active for example on ATP and 4-nitrophenyl-dTMP, and to be a Zn^2+^ ectometalloenzyme which is inactivated by EDTA (Bischoff et al. 1975; Cameselle et al. 1984; López-Gómez et al. 1998; Ribeiro et al. 2000; Stefan et al. 1997). While studying RLNPP/PDE inhibition by a culture-grade preparation of acidic fibroblast growth factor (FGF-1), it was found that the inhibition was due to the presence of contaminant EDTA (López-Gómez et al. 1998; Stefan et al. 1997). The inhibition was time-dependent, blocked by the substrate 4-nitrophenyl-dTMP, and potentiated (accelerated) by glycine used as buffer instead of Tris, which was attributed to a conformational change evoked by the amino acid (López-Gómez et al. 1998). This old finding has not been further extended, except for a PhD thesis that remains unpublished (Romero 2003).

The time-dependent RLNPP/PDE inactivation is here treated as a first-order reaction with respect to active enzyme, considering [EDTA] invariant, enabling quantification in terms of apparent inactivation constants (k_i(ap)_). This was studied with all the common amino acids that do not affect RLNPP/PDE activity by themselves, i.e. all except cysteine and histidine. The acceleration factors showed, with some conspicuous exceptions, good correlations with the stability constant of the Zn^2+^-amino acid complexes (positive correlation) and with amino acid size (negative correlation). The results are compatible with a conformational change evoked by direct interaction of free amino acids with RLNPP/PDE.

Free amino acids are relevant to insulin activity and resistance (Chen et al. 2022; Gar et al. 2018; Krebs et al. 2002; Lee et al. 2018; Patti et al. 1998; Saleem et al. 2019; Seibert et al. 2015; Tremblay et al. 2007), what could eventually be related to the role of human ENPP1 as negative modulator of the insulin receptor (Grarup et al. 2006; Maddux et al. 2006; Maddux et al. 1995; Tassone et al. 2018; Zhou et al. 2009). For this reason, experiments were designed to find out what rat enzymes are present in RLNPP/PDE. This included the assay of hydrolytic activity on the STING (stimulator of interferon genes) agonist 2’,3’-cGAMP (2’,3’-cyclic-GMP-AMP), a feature of human ENPP1 (Borza et al. 2021; Kato et al. 2018; Li et al. 2014; Ritchie et al. 2022) and ENPP3 (Mardjuki et al. 2024). Data obtained by LC-MS/MS (liquid chromatography-tandem mass spectrometry) indicated that RLNPP/PDE did not contain Enpp1, but Enpp3, thus confirming the 2’,3’-cGAMP hydrolase activity of the latter. The significance of the effect of amino acids on Enpp3 is discussed in the light of the known functions of human ENPP3 and orthologs.

## Materials and Methods

### Chemicals, biochemicals and chromatography media

The 20 common L-amino acids (collection LAA-21), p-aminobenzoic acid, γ-aminobutyric acid, ε-aminohexanoic acid, D-aspartate, n-butyric acid, glutaric acid, L-lactic acid, malonic acid, oxaloacetic acid, propionic acid, D-alanine, β-alanine, 2-aminobutane, aminoethane, formamide, trans-4-hydroxy-L-proline, 1-aminopropane, 2-aminopropane, ethylene glycol, glycylglycine, glycylglycylglycine, 4-nitrophenyl-dTMP sodium salt, 4-nitrophenyl-phosphorylcholine and Triton X-100 were from Sigma (now Merck Life Sciences, Madrid, Spain). Ammonium acetate and aminomethane were from Aldrich (now Merck Life Sciences, Madrid, Spain). L-Malic acid, succinic acid, ammonium chloride, urea, hydrochloric acid, sodium chloride, magnesium chloride, sodium acetate, monosodium phosphate, bisodium phosphate, EDTA bisodium salt (Titriplex-III) and sodium hydroxide were from Merck (now Merck Life Sciences, Madrid, Spain). α-Ketoglutarate, pyruvate, bovine liver catalase, bovine pancreas trypsin, soybean trypsin inhibitor, bovine serum albumin, sucrose and Tris were from Boehringer/Roche (now Merck Life Sciences, Madrid, Spain). 2’,3’-cyclic-GMP-AMP was from Biolog Life Science Institute, Bremen, Germany. Sephacryl S-200 and Sephadex G-25 (PD-10 columns) were from Amer-sham-Pharmacia (Cytiva, now purchasable from Merck Life Sciences, Madrid, Spain). DEAE-cellulose DE52 was from Whatman (now purchasable from Merck Life Sciences, Madrid, Spain).

### Preparation of crude rat liver membranes

Female Wistar rats of about 250 g, fed ad libitum with commercial pellet food and water, were used and euthanized. Fresh liver was homogenized in 0.25 M sucrose (3 mL per g of tissue) with a Potter-Elvehjem homogenizer (with ground glass pestle and tube) cooled on ice. The homogenate was centrifuged for 2 min at 600 g and the supernatant collected was centrifuged for 60 min at 100,000 g. The supernatant was discarded and the pre-cipitate was resuspended in 20 mM Tris-HCl pH 8.7. This crude preparation was used to test the response of membrane RLNPP/PDE to incubation with EDTA in the absence and presence of glycine.

### Solubilization of RLNPP/PDE

The enzyme was solubilized from crude rat liver membranes, except that the last precipitate was resuspended in 20 mM Tris-HCl, pH 8.7, supplemented with 5 mM MgCl_2_ and 10 mg Triton X-100/mL. After standing 30 min at 4°C, the resuspended precipitate was centrifuged for 60 min at 100,000 g and the supernatant, containing 10–15 mg protein/mL, was treated by limited trypsinization at 37°C with a total of 31 µg of trypsin per mg of protein dosed as follows: 12,5 µg/mg were added immediately and 6,25 µg were added after 3.5, 6.5 and 7.5 h, respectively. The trypsinization was stopped 8.5 h after starting the treatment by addition of 62 µg of soybean trypsin inhibitor per mg of protein treated. The resulting sample was then dialyzed for 12 h at 4°C against 50 volumes of Tris-HCl, pH 8.7, with 5 mM MgCl_2_, and afterwards centrifuged for 60 min at 100,000 g.

### Partial purification of solubilized RLNPP/PDE

The solubilized preparation was adsorbed to a DEAE-cellulose column of 25 cm x 1.5 cm equilibrated in Tris-HCl, pH 8.7, with 5 mM MgCl_2_. The activity assayed with 4-nitrophenyl-dTMP as substrate was recovered with a linear 0–400 mM NaCl gradient in the same buffer. A single activity peak was collected around 130 mM NaCl, concentrated by ultrafiltration through an Amicon PM30 membrane under nitrogen pressure, and applied to a Sephacryl S-200 column of 88 cm x 1.7 cm equilibrated in and eluted with 20 mM Tris-HCl, pH 8.7, with 5 mM MgCl_2_. A single peak of activity was recovered with V_e_ 101 mL. It was concentrated again by ultrafiltration and was submitted to buffer exchange in a Sephadex G-25 column of 5 cm x 1.4 cm equilibrated in 5 mM sodium phosphate pH 8.25. It was supplemented with 1 mg bovine serum albumin/mL and divided in portions that were frozen at -20°C. Different preparations showed activities of ≈2–6 units/mL. The same procedure was followed to obtain the RLNPP/PDE preparation used in other studies (López-Gómez et al. 1998; Ribeiro et al. 2000).

### RLNPP/PDE activity assay

The standard RLNPP/PDE activity assay was run discontinuously measuring the formation of nitrophenol from 1 mM 4-nitrophenyl-dTMP in 50 mM Tris-HCl, pH 9.0, at 37°C. After different incubation lengths, the reaction was finished by adding 1 mL of 0.2 M NaOH over 0.2-mL reaction mixtures or over 0.2-ml aliquots taken from larger reaction mixtures. The amount of nitrophenol was determined from A_405_ measurements (ε = 18,500 M^-1^cm^-1^) which correspond to the absorbance produced by anion nitrophenolate, since nitrophenol has a pK_a_ 7.14. To study the activity of RLNPP/PDE over 2’,3’-cGAMP, reaction mixtures at 37°C contained in 100 µL 50 mM Tris-HCl, pH 9, 10 µM 2’,3’-cGAMP and 1 µL or 10 µL of enzyme. Under assay conditions, the activity was proportional to the amount of RLNPP/PDE and linear with incubation length.

### HPLC analysis of RLNPP/PDE action on 2’,3’-cGAMP

HPLC was performed essentially as described (Ribeiro et al. 2023) on a Tracer Excel 120 column (150 mm × 4 mm) protected by a pre-column (10 mm × 4 mm) of the same material (octadecylsilica; Teknokroma, San Cugat del Vallés, Barcelona). An HP1100 system was used adjusted to measure A_260_. Samples of 20 µl were injected and the elution was performed at 0.5 ml/min with two buffers: A, 5 mM Na-phosphate, pH 7.0, 5 mM tetrabutylammonium, 20% methanol (by vol.); B, 100 mM Na-phosphate, pH 7.0, 5 mM tetrabutylammonium, 20% methanol. The initial mobile phase was 100% A, and a linear gradient was applied up to 50% A and 50% B in 4 min, followed by another linear gradient up to 100% B in 1 min and isocratic elution with 100% B for 5 min. Retention times (min) were: Guo (3.9), Ado and GMP (5.2), AMP (5.9) and 2’,3’-cGAMP (7.3).

### RLNPP/PDE inactivation by EDTA

Enzyme inactivation by EDTA was studied in the absence of added bivalent cations. Under standard conditions, RLNPP/PDE treatment was performed in mixtures containing 50 mM Tris, pH 9.0, 5 µM EDTA and one of the products tested as inactivation modifiers (amino acids or analogs). Controls were carried out without any modifier. These mixtures were preincubated for 10 min at 37°C, and the experiments were initiated by the addition of RLNPP/PDE and terminated by addition of 1 mM 4-nitrophenyl-dTMP. This last addition served a twofold purpose: it blocked the inactivation and started the assay of residual RLNPP/PDE activity. Usually, to study the kinetics of RLNPP/PDE inactivation, increasing periods of inactivation were carried out in a single mixture from which 196 µL aliquots (or multiples) were removed at defined times and added over tubes containing 4 µl (or the respective multiples) of 50 mM 4-nitrophenyl-dTMP. This started the assay of residual RLNPP/PDE activity. The assays were run in triplicate.

The kinetics of inactivation of RLNPP/PDE by EDTA can be described according to the binary reaction E-Zn + EDTA → E + EDTA-Zn, either by direct trapping of enzyme-bound metal or after slow metal release followed by rapid chelation. The apparent second-order equation for the whole process is

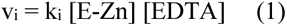

which can be simplified to an apparent first-order equation considering that under the experimental conditions [EDTA] >> [E-Zn]. Therefore [EDTA] remains approximately constant during the inactivation allowing to define an apparent k_i_ as

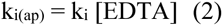

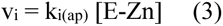

The time courses of inactivation by several EDTA concentrations are represented as direct plots of residual RLNPP/PDE activity versus time and as semilogarithmic transformations of the same data. The slope of linear semilogarithmic plots equals –k_i(ap)_. The k_i(ap)_ value increases linearly with EDTA concentration. If desired, k_i(ap)_ can be divided by [EDTA] to obtain an EDTA-independent k_i_ value, However, we have preferred to use k_i(ap)_ values obtained at indicated EDTA concentrations (most frequently 5 µM EDTA).

The Pearson coefficients and *P* values for the correlation between the increase of k_i(ap)_ and different parameters were calculated with the Pearson Correlation Coefficient Calculator in the web page Social Science Statistics (https://www.socscistatistics.com/, accessed on 27 June, 2024).

### Preparation of modifiers of the inactivation of RLNPP/PDE by EDTA

Amino acids and analogs were dissolved individually at 100 mM concentration and adjusted at pH 9.0 by addition of HCl or NaOH as needed. Due to solubility limits, cysteine was dissolved at 20 mM and tyrosine at 6.25 mM. The latter was dissolved at 45°C immediately before addition to reaction mixtures.

Amino acid mixtures were prepared from a stock solution at pH 9.0 in which each amino acid was at a concentration 20-fold that prevailing under fasting conditions in rat portal blood (Patti et al. 1998) (see Table 1).

**Table 1.** Data for amino acids included in mixtures to be tested for their effect on the inactivation of RLNPP/PDE by EDTA. The acceleration factor (AF) equals the ratio of the k_i(ap)_ obtained in the presence of the individual amino acid versus that in its absence (Fig. 3). The k_i(ap)_ increment was calculated as 100x(AF–1). Amino acid levels in rat portal blood were taken from (Patti et al. 1998). The fraction of effect in a 5 mM amino acid mixture was calculated by multiplying the k_i(ap)_ increase by the molar fraction of the amino acid in the mixture at fasting concentrations

| Amino acid<br>(5 mM) | $k_{i(ap)}$<br>($\text{min}^{-1}$)<sup>1</sup> | Acceleration<br>factor (AF) | $k_{i(ap)}$<br>increase<br>($100[AF-1]$)<br>(%) | Amino acid<br>concentration in rat<br>portal blood under<br>fasting<br>($\mu\text{M}$) | $k_{i(ap)}$ increase produced by<br>each amino acid<br>in a 5 mM amino acid<br>mixture<br>(%) |
| --- | --- | --- | --- | --- | --- |
| None | 0.054 | 1.0 | 0 | — | — |
| Glycine | 0.355 | 6.6 | 560 | 300 | 58.23 |
| Tryptophan | 0.333 | 6.2 | 520 | 70 | 12.62 |
| Aspartate | 0.331 | 5.3 | 430 | 30 | 4.47 |
| Serine | 0.230 | 4.4 | 340 | 200 | 23.57 |
| Threonine | 0.196 | 3.9 | 290 | 180 | 18.09 |
| Asparagine | 0.189 | 3.7 | 270 | 60 | 5.62 |
| Alanine | 0.180 | 3.7 | 270 | 400 | 37.44 |
| Glutamate | 0.172 | 3.4 | 240 | 100 | 8.32 |
| Tyrosine | 0.217 | 3.2 | 220 | 75 | 5.72 |
| Phenylalanine | 0.170 | 3.1 | 210 | 50 | 3.64 |
| Methionine | 0.151 | 2.7 | 170 | 40 | 2.36 |
| Glutamine | 0.130 | 2.7 | 170 | 350 | 20.62 |
| Leucine | 0.114 | 2.1 | 110 | 250 | 9.53 |
| Arginine | 0.102 | 2.0 | 100 | 100 | 3.47 |
| Valine | 0.115 | 2.0 | 100 | 180 | 6.24 |
| Isoleucine | 0.104 | 1.8 | 80 | 100 | 2.77 |
| Lysine | 0.093 | 1.7 | 70 | 300 | 7.28 |
| Proline | 0.089 | 1.6 | 60 | 100 | 2.08 |
| Total |  |  |  | 2945 | 232.06 |

<sup>1</sup> The values of $k_{i(ap)}$ were obtained from inactivation experiments carried out in triplicate (Fig. 3). The $k_{i(ap)}$ in the absence of amino acid is the mean obtained in all the experiments of Fig. 3

### Proteomic analysis by LC-MS/MS

In summary, A 0.4-mL concentrated sample of RLNPP/PDE from the Sephacryl S-200 step containing 0.15 mg protein/mL was concentrated tenfold by speed-vac. The proteins were precipitated in chloroform/methanol at 4°C, resuspended in 50 mM ammonium bicarbonate, disulfide linkages were reduced with 10 mM dithiothreitol and blocked with 22.5 mM iodoacetamide. Proteolysis was run overnight at 37°C with 1:50 (w/w) recombinant trypsin (Roche). The peptides obtained were fractionated in a 44-min chromatographic run (Evosep One, Evosep) followed by electrospray ionization and analysis in a TIMS-TOF Pro 2 spectrometer (Bruker) set in DDA (da-ta-dependent acquisition) mode. Precursor peptides were fragmented by high collision energy dissociation and the MS/MS spectra were collected and processed off-line with the MSFragger/FragPipe software (Kong et al. 2017) against the Rattus norvegicus proteome database (Uniprot/Swissprot). Sample treatment and analysis were performed in the Proteomic Unit (CAI Biological Techniques) of Complutense University of Madrid (Madrid, Spain), who provided the list of identified proteins (Table S1).

## Results

### Quantification of the inactivation of RLNPP/PDE by EDTA

The course of the time-dependent inactivation of RLNPP/PDE at different EDTA concentrations is shown in Fig. 1A. In the absence of EDTA, the activity was stable over the assay period (60 min), whereas EDTA at 1 µM, 5 µM and 30 µM evoked progressively faster inactivation. The corresponding semilogarithmic plots are shown in Fig. 1B, from where k_i(ap)_ values were derived from the slopes of the linear parts of the plots (k_i(ap)_ = -slope). The insert of Fig. 1B shows that k_i(ap)_ was directly proportional to EDTA concentration.

**Fig. 1.**
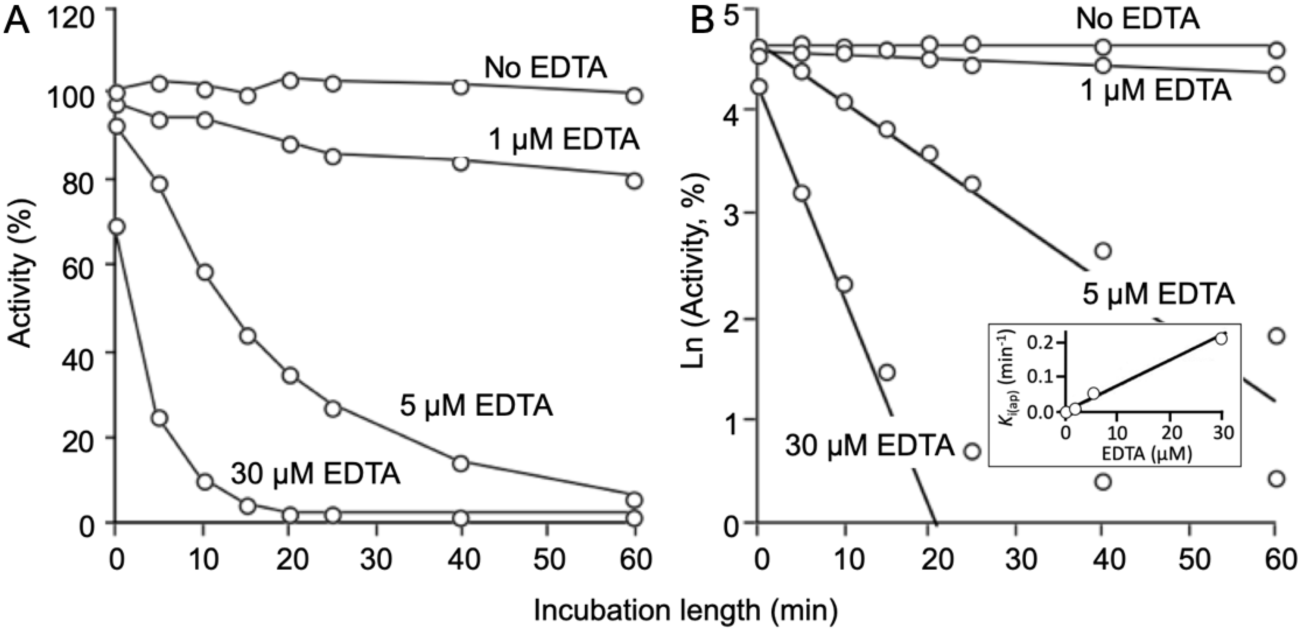
Dependence of RLNPP/PDE inactivation on EDTA concentration (A) and estimation of k_i(ap)_ at different EDTA concentrations (B). The values of k_i(ap)_ are plotted in the insert versus EDTA concentration

### Effect of common amino acids on the inactivation of RLNPP/PDE by EDTA

The effect of glycine on the inactivation of RLNPP/PDE by EDTA was tested at various amino acid concentrations. In 5-min incubations with 6 µM EDTA, in the presence of 10–50 mM glycine, the inactivation of the enzyme was practically 100%, while in the presence of 1–5 mM glycine, 75–90% inactivation was observed (Fig. 2).

**Fig. 2.**
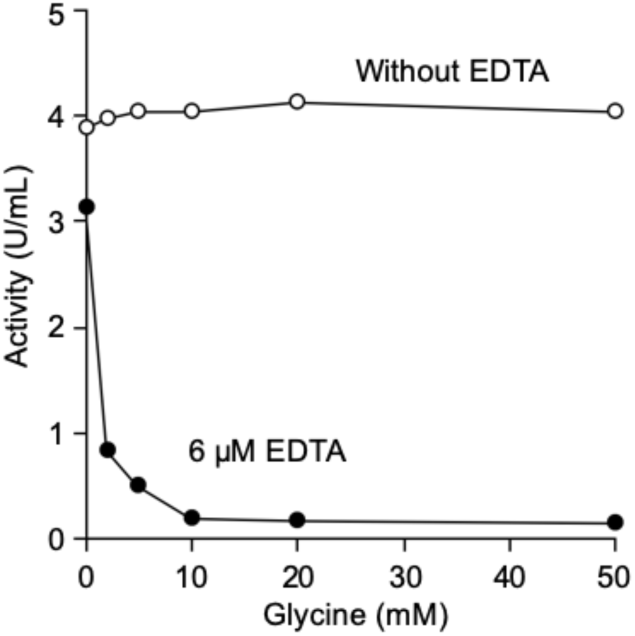
Inactivation of RLNPP/PDE by 5-min incubation with EDTA in the presence of increasing glycine concentration

After the preliminary tests shown in Fig. 1 and Fig. 2, the selected conditions to study the effect of common amino acids on k_i(ap)_ were incubation with 5 µM EDTA and 5 mM amino acid. The twenty common amino acids plus hydroxyproline were studied: nineteen of them did not affect RLNPP/PDE activity in the absence of EDTA but accelerated enzyme inactivation by EDTA (Fig. 3). The numerical data are summarized in Table 1, including values of AF (acceleration factor) and of percent increase of k_i(ap)_.

**Fig. 3.**
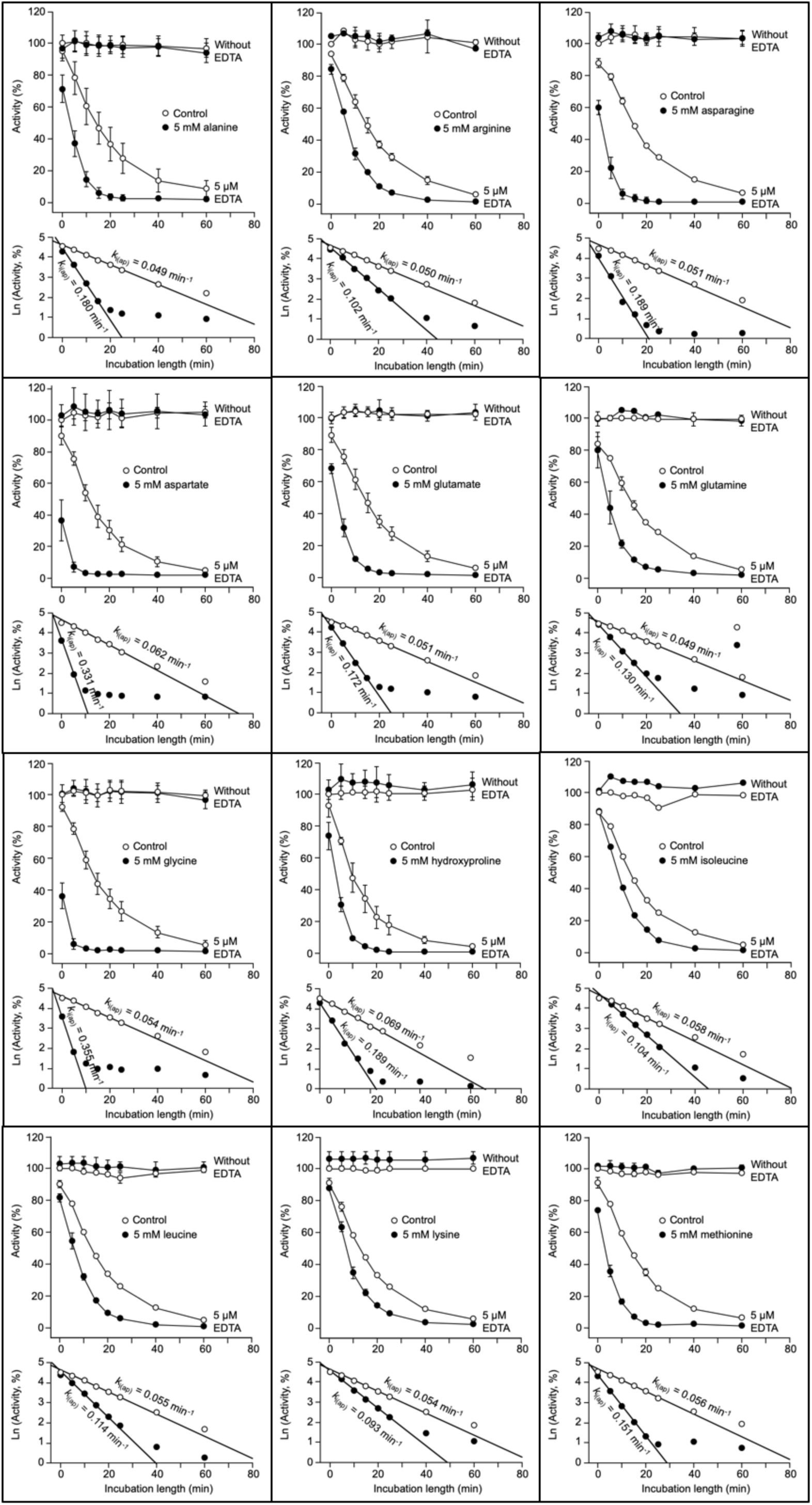
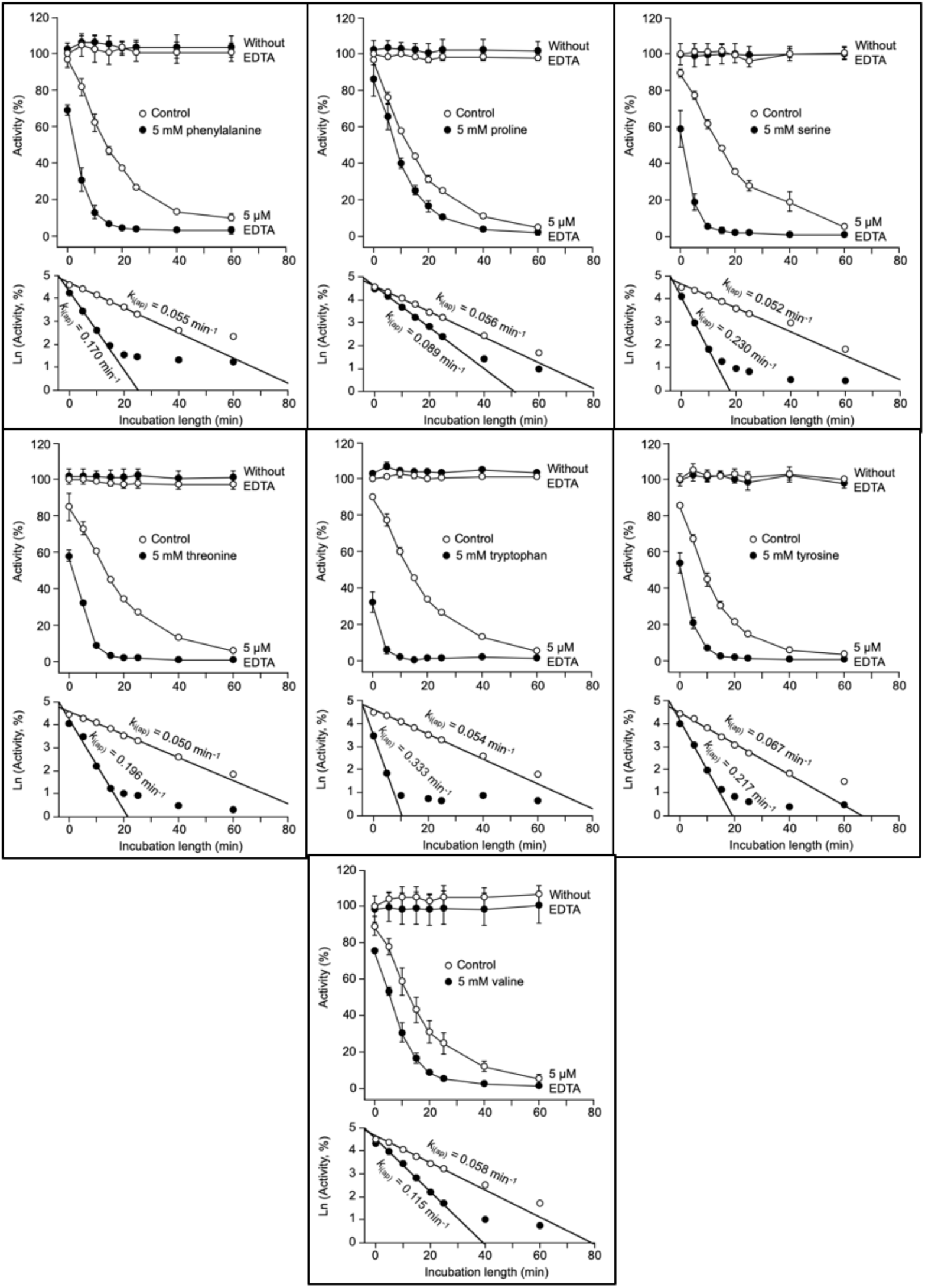
Acceleration of the time-dependent inactivation of RLNPP/PDE by EDTA: effects of nineteen common amino acids that do not affect enzyme activity in the absence of EDTA. The course of inactivation and semilogarithmic plots are shown for each amino acid. Activity assays were performed in triplicate. Error bars are standard deviations

In contrast to those amino acids, cysteine and histidine inhibited RLNPP/PDE strongly in the absence of EDTA, what difficulted the quantification of the acceleration evoked by these amino acids, since in the presence of EDTA and amino acid a full inhibition was recorded (Fig. 4).

**Fig. 4.**
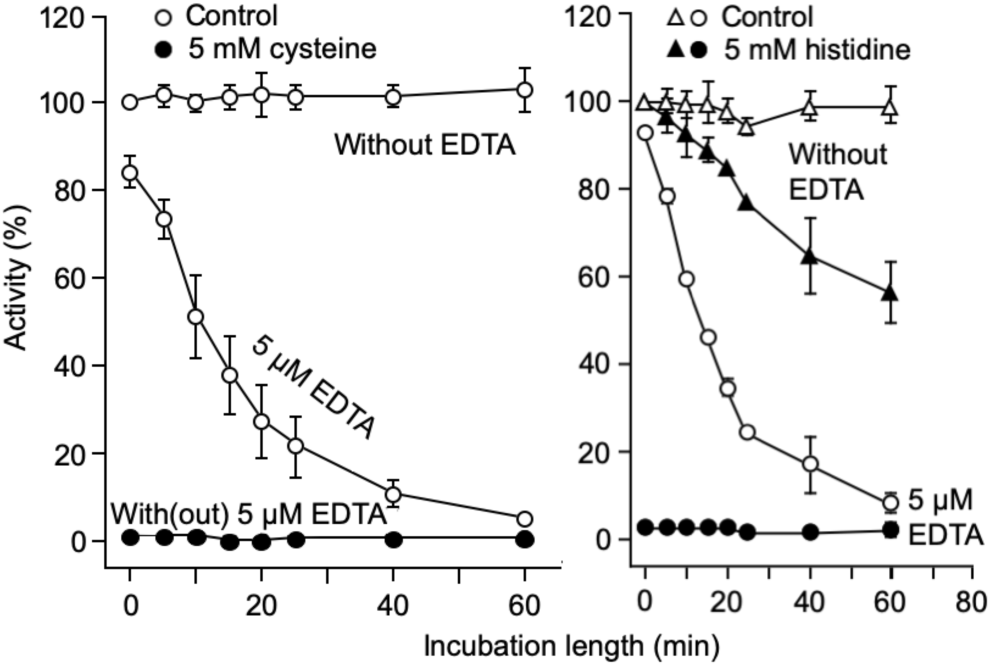
Acceleration of the time-dependent inactivation of RLNPP/PDE by EDTA: effects of two common amino acids that are strong inhibitors of enzyme activity in the absence of EDTA

### Effect of amino acid mixtures on the inactivation of RLNPP/PDE by EDTA

Common amino acid mixtures were prepared at different concentrations based on those prevailing in rat portal blood during fasting (Patti et al. 1998). Histidine and cysteine were omitted in these mixtures because their strong inhibitory effect in the absence of EDTA precludes a clear estimation of their effect on RLNPP/PDE inactivation by EDTA (Fig. 3). The fasting concentrations of individual amino acids range from 30–400 µM, yielding a total concentration of 2,945 µM (Table 1). Using the results obtained with the individual amino acids at 5 mM concentration (Fig. 3), acceleration factors were calculated and converted to percent k_i(ap)_ increases. Multiplying these values by the molar fraction of the corresponding amino acid in the fasting mix, one obtains the fraction of effect produced by each amino acid in a mix where the total concentration were 5 mM and the amino acid proportions were like in fasting conditions. The sum of these values provides a theoretical prediction of the effect expected for such a 5 mM mixture (Table 1). Fig. 5 shows the effects produced by amino acid mixtures at different total concentrations, ranging from 0.25–4 fold the concentration under fasting. The effect of the 5 mM mixture of amino acids (1.7 fold the fasting concentration) was in good agreement with the theoretical prediction (see the insert of Fig. 5).

**Fig. 5.**
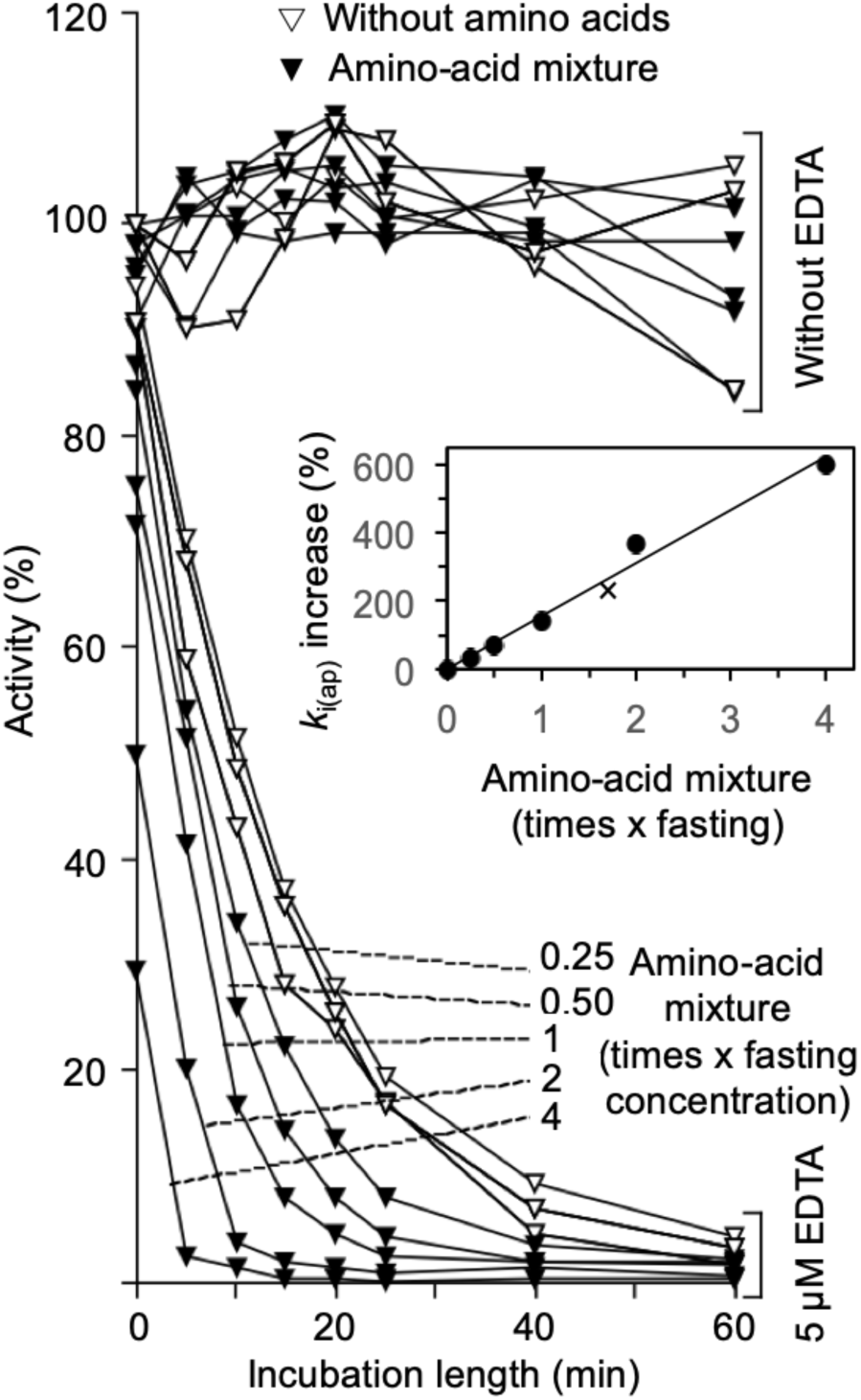
Acceleration of the time-dependent inactivation of RLNPP/PDE by EDTA: amino acid mixtures of different concentration. The composition of the mixtures is shown in Table 1. The amino acid concentrations were 0.25–4 fold those prevailing under fasting conditions in rat portal blood. The insert shows the proportionality between the concentration of the mixture and the k_i(ap)_ increase; the X symbol marks the theoretical prediction made in Table 1 for a 5 mM mixture (1.7-fold the fasting concentration)

### Effect of amino acid analogs on the inactivation of RLNPP/PDE by EDTA

All the common amino acids studied accelerated RLNPP/PDE inactivation by EDTA to different degrees, with the only apparent exception of cysteine. In this case, the strong inhibition caused at 5 mM concentration in the absence of EDTA, precluded to observe any effect in the presence of EDTA (Fig. 3). In the case of histidine, which in the absence of EDTA was itself a significant time-dependent inhibitor of RLNPP/PDE, the inactivation by 5 µM EDTA was potentiated indeed, but with both inhibitors together (histidine and EDTA) the time-dependency was not apparent (Fig. 4). With the rest of common amino acids, the increase of k_i(ap)_ ranged from 60% by proline to >500% by glycine and tryptophan (Table 1). This variation gives information about the role of the amino acid side chain in the interaction with RLNPP/PDE (see the Discussion), but not about the rest of the amino acid structure (—CHNH_2_—COO^-^). To investigate this question, analogs and structural variants of common amino acids were tested, including two D-amino acids (D-aspartate and D-alanine), analogs devoid of the α-amine group (acetate, propionate, malonate, butyrate, succinate and glutarate), α-oxoacids (pyruvate, oxaloacetate and α-oxoglutarate), α-hydroxyacids (L-lactate and L-malate), analogs devoid of the α-carboxyl group (aminomethane, aminoethane, 1-aminopropane and 2-aminopropane, 2-aminobutane), non-α-amino acids with increasing distances between the carboxyl and the amine groups (β-alanine, γ-aminobutyrate and δ-aminohexanoate), glycine peptides which also increase the distance between the carboxyl and the amine groups (glycylglycine and glycylglycylglycine), and other miscellaneous compounds (ammonium chloride, ethylene glycol, formamide, urea, p-aminobenzoate). The results obtained with all these variants are shown in Table 2 and discussed later.

**Table 2.** Effects of amino acid analogs on the inactivation of RLNPP/PDE by EDTA.

| Amino acid analog<br>(5 mM) | Acceleration<br>factor (AF) | $k_{i(ap)}$ increase<br>(100[AF-1])<br>(%) |
| --- | --- | --- |
| D-Aspartate | 5.3 | 430 |
| D-Alanine | 2.3 | 130 |
| Pyruvate | 1.6 | 60 |
| Glycylglycine | 1.5 | 50 |
| α-Ketoglutarate | 1.4 | 40 |
| Ammonium acetate | 1.3 | 30 |
| Aminoethane | 1.2 | 20 |
| Aminomethane | 1.2 | 20 |
| 2-Aminopropane | 1.2 | 20 |
| Butyrate | 1.2 | 20 |
| Ammonium chloride | 1.2 | 20 |
| Glycylglycylglycine | 1.2 | 20 |
| L-Lactate | 1.2 | 20 |
| Malonate | 1.2 | 20 |
| Oxaloacetate | 1.2 | 20 |
| 2-Aminobutane | 1.1 | 10 |
| ε-Aminohexanoate | 1.1 | 10 |
| Glutarate | 1.1 | 10 |
| L-Malate | 1.1 | 10 |
| Propionate | 1.1 | 10 |
| Succinate | 1.1 | 10 |
| Sodium acetate | 1.0 | 0 |
| γ-Aminobutyrate | 1.0 | 0 |
| 1-Aminopropane | 1.0 | 0 |
| Ethylene glycol | 1.0 | 0 |
| Formamide | 1.0 | 0 |
| Urea | 1.0 | 0 |
| β-Alanine | 0.9 | -10 |
| p-Aminobenzoate | 0.8 | -20 |

### Effect of glycine on the inactivation of RLNPP/PDE by EDTA in crude membrane preparations

In previous sections, the effect of amino acids on RLNPP/PDE inactivation was studied with a solubilized enzyme preparation. However, in its natural location the enzyme is bound to membranes. Fig. 6 shows the results of testing the effect of 5 mM glycine on the inactivation of the RLNPP/PDE activity by 10 µM EDTA in a crude preparation of rat liver membranes. The effect was tested on freshly obtained membranes and in membranes kept at 4°C for 2-6 days. The value of k_i(ap)_ in the absence of glycine did not change with time elapsed. In contrast, the accelerating effect of glycine, that was clear in every case, became stronger in aged membranes, approaching the values observed with solubilized RLNPP/PDE. This result argues strongly against the formation of EDTA-Zn^2+^-amino acid complexes in solution as possible explanation for the pro-inactivating effect of amino acids (see the Discussion).

**Fig. 6.**
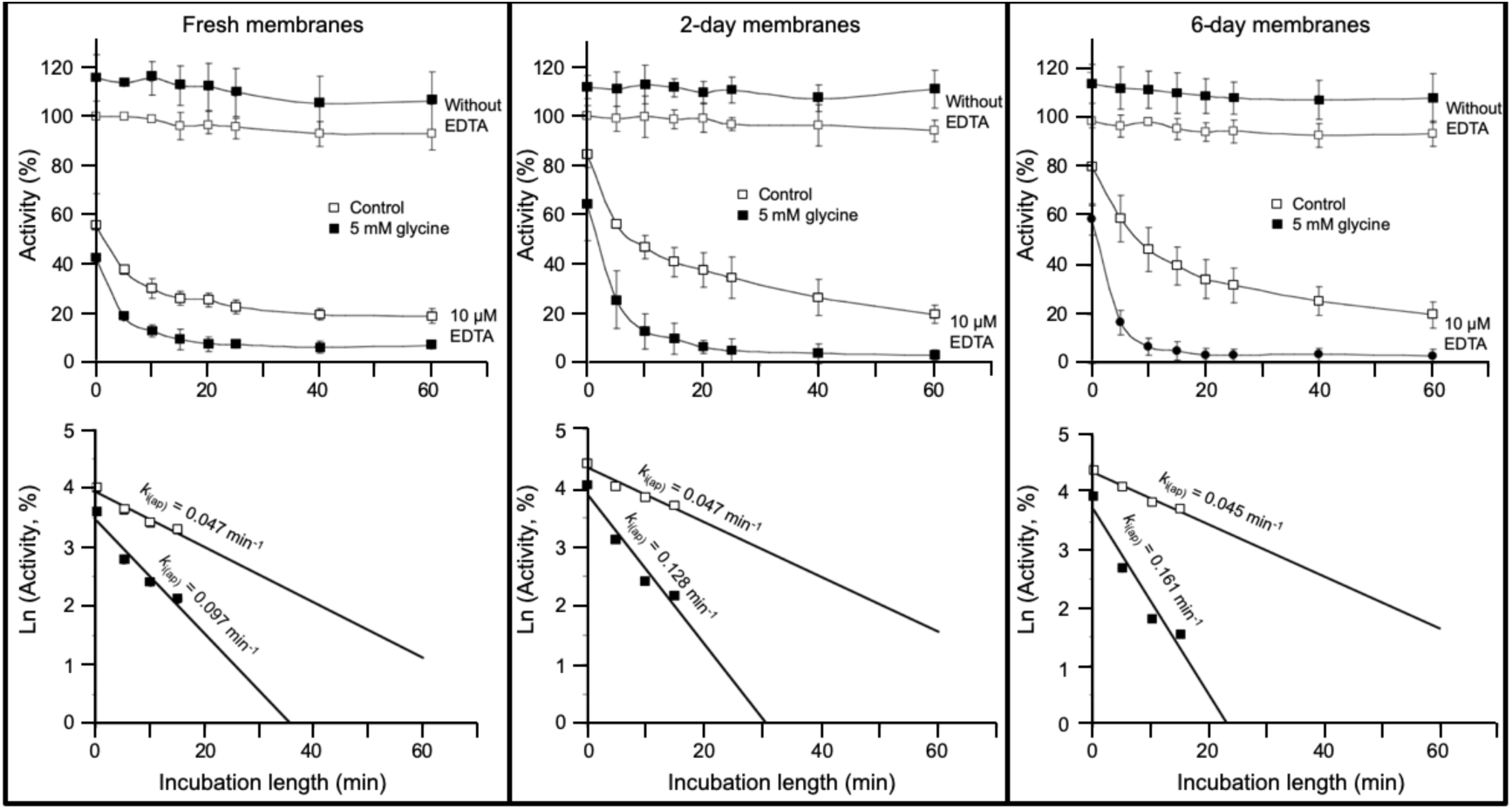
Acceleration by glycine of the time-dependent inactivation of RLNPP/PDE by EDTA in a crude preparation of liver membranes. With fresh membranes, the experiment was started immediately after preparation and finished in less than two hours. In the other cases, the membranes were kept at 4°C for two or six days before the experiments

### Experiments designed to find out the molecular identity of the enzyme responsible for RLNPP/PDE activities: assays of the hydrolysis of 2’,3’-cGAMP, ATP and 4-nitrophenylphosphorylcholine, and proteomic analysis by LC-MS/MS

The hydrolysis of 2’,3’-cGAMP is a well-known activity of human ENPP1 (Borza et al. 2021; Kato et al. 2018; Li et al. 2014; Ritchie et al. 2022). However, while this work was in progress, this activity was also described for human SMPDL3A (ASM3A_HUMAN, acid sphingomyelinase-like phosphodiesterase 3a) (Hou et al. 2023) and for human ENPP3 (Mardjuki et al. 2024). Like RLNPP/PDE, these three proteins are also active against ATP and 4-nitrophenyl-dTMP. Therefore, the 2’,3’-cGAMP hydrolase activity of RLNPP/PDE was assayed by HPLC with positive results (Fig. 7), pointing to the possible presence of rat orthologs of any of those enzymes. As a complement to this study, the activity of RLNPP/PDE on ATP was also assayed by HPLC (Fig. 7), and that on 4-nitrophenyl-phosphorylcholine (typical of SMPDL3A) was assayed by A_405_ measurements with detection of an activity much smaller (<1%) than that on 4-nitrophenyl-dTMP (Table 3).

**Fig. 7.**
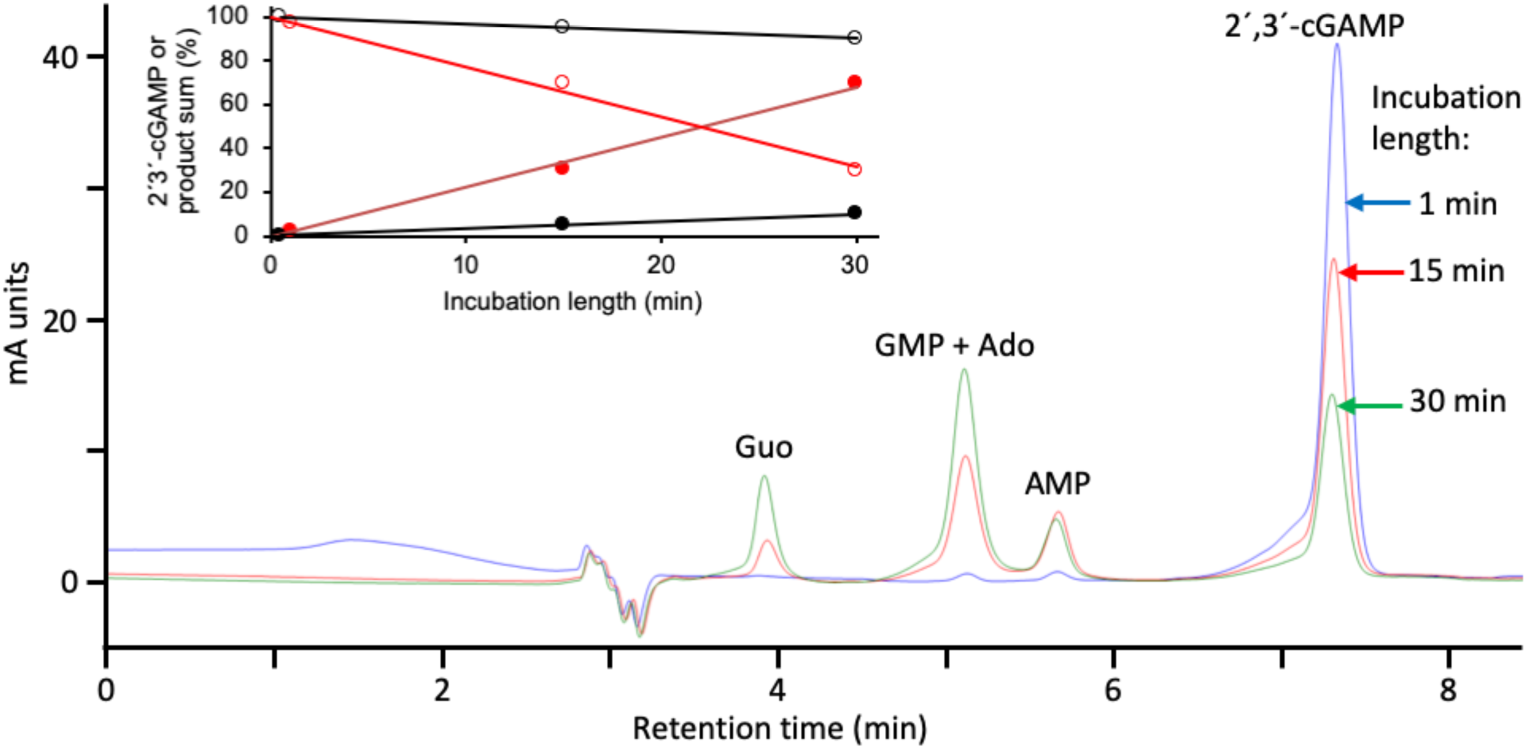
RLNPP/PDE hydrolysis of 2’,3’-cGAMP. A reaction mixture of 100 µL containing 50 mM Tris-HCl, pH 9, 10 µM 2’,3’-cGAMP and 10 µL of RLNPP/PDE was incubated at 37°C. At the indicated times, 20 µL were analyzed by HPLC. The insert shows the quantitative results of reaction mixtures with either 1 µL (black line and symbols) or 10 µL (red line and symbols) of enzyme. The data show the remaining substrate (open circles) and the sum of AMP, GMP, Ado and Guo formed (closed circles). 5’-Nucleotides were direct products of RLNPP/PDE and nucleosides were secondary products due to a contaminating nucleotidase activity

**Table 3.** RLNPP/PDE hydrolytic activities.

| Substrate | Activity<sup>1</sup><br>(nmol min<sup>-1</sup> mL<sup>-1</sup>) |
| --- | --- |
| 4-Nitrophenyl-dTMP (20 mM) | 1836 ± 238 |
| 4-Nitrophenyl-phosphorylcholine (20 mM) | 15 ± 7 |
| ATP (100 μM) | 157 ± 17 |
| 2',3'-cGAMP (10 μM) | 2.7 ± 0.6 |

<sup>1</sup> Activities were assayed at pH 9 and 37°C. The hydrolysis of the 4-nitrophenyl derivatives was assayed by measuring the formation of nitrophenol at 405 nm. The hydrolysis of 2',3'-cGAMP and ATP were assayed by HPLC (Fig. 6 and Fig. 7)

The proteomic analysis of RLNPP/PDE by LC-MS/MS (Table S1) revealed the absence of four members of the ENPP family (Enpp1, Enpp2, Enpp6 and Enpp7), and the presence of other three (Enpp3, Enpp4, Enpp5). In addition, the presence of Smpdl3a was also detected (Table 4). Enpp4 and Enpp5 are orthologs of the murine and human proteins Enpp4/ENPP4 and Enpp5/ENPP5. Smpdl3a is the ortholog of murine and human proteins Smpdl3a/SMPDL3A. Anyhow, it should be considered that detection of peptides of a specific enzyme by LC-MS/MS does not necessarily mean that active protein was present in the sample. In the Discussion, all these enzyme-activity and mass-spectrometry data are used in support of the attribution of RLNPP/PDE activities to rat Enpp3.

**Table 4.** Selected proteins identified in RLNPP/PDE by LC-MS/MS. Data taken from Table S1, except gene names that were taken from the Uniprot database.

| Uniprot name | Description | Coverage % | Total peptides | Total spectral count | Spectral abundance factor | Gene name |
| --- | --- | --- | --- | --- | --- | --- |
| ENPP3_RAT | Ectonucleotide pyrophosphatase /phosphodiesterase family member 3 | 15.31 | 12 | 23 | 0.026 | Enpp3 |
| A6JJ18_RAT | bis(5'-adenosyl)-triphosphatase | 30.84 | 6 | 13 | 0.029 | Enpp4 |
| ENPP5_RAT | Ectonucleotide pyrophosphatase /phosphodiesterase family member 5 | 19.08 | 5 | 6 | 0.013 | Enpp5 |
| ASM3A_RAT | Acid sphingomyelinase-like phosphodiesterase 3a | 8.31 | 2 | 5 | 0.011 | Smpdl3a |

## Discussion

### The evidence favors Enpp3 as responsible for the enzyme activities of RLNPP/PDE and for the acceleration of the EDTA-dependent inactivation by amino acids

For this discussion, one has to consider first all the enzyme activities shown by the RLNPP/PDE preparation in this study (Table 3) and in previous ones, i.e. phosphodiesterase activity on 4-nitrophenyl-dTMP (Table 3 and (López-Gómez et al. 1998; Ribeiro et al. 2000)) and 4-nitrophenyl-phosphorylcholine (Table 3), nucleotide pyrophosphatase activity on ATP (Fig. 7, Table 3 and (Ribeiro et al. 2000)) and 2’,3’-cGAMP hydrolase (Fig. 6 and Table 3). On the other hand, these activities should be compared to those of the relevant candidates, namely Enpp3, Enpp4, Enpp5 and Smpdl3a (Table 4).

**Fig. 8.**
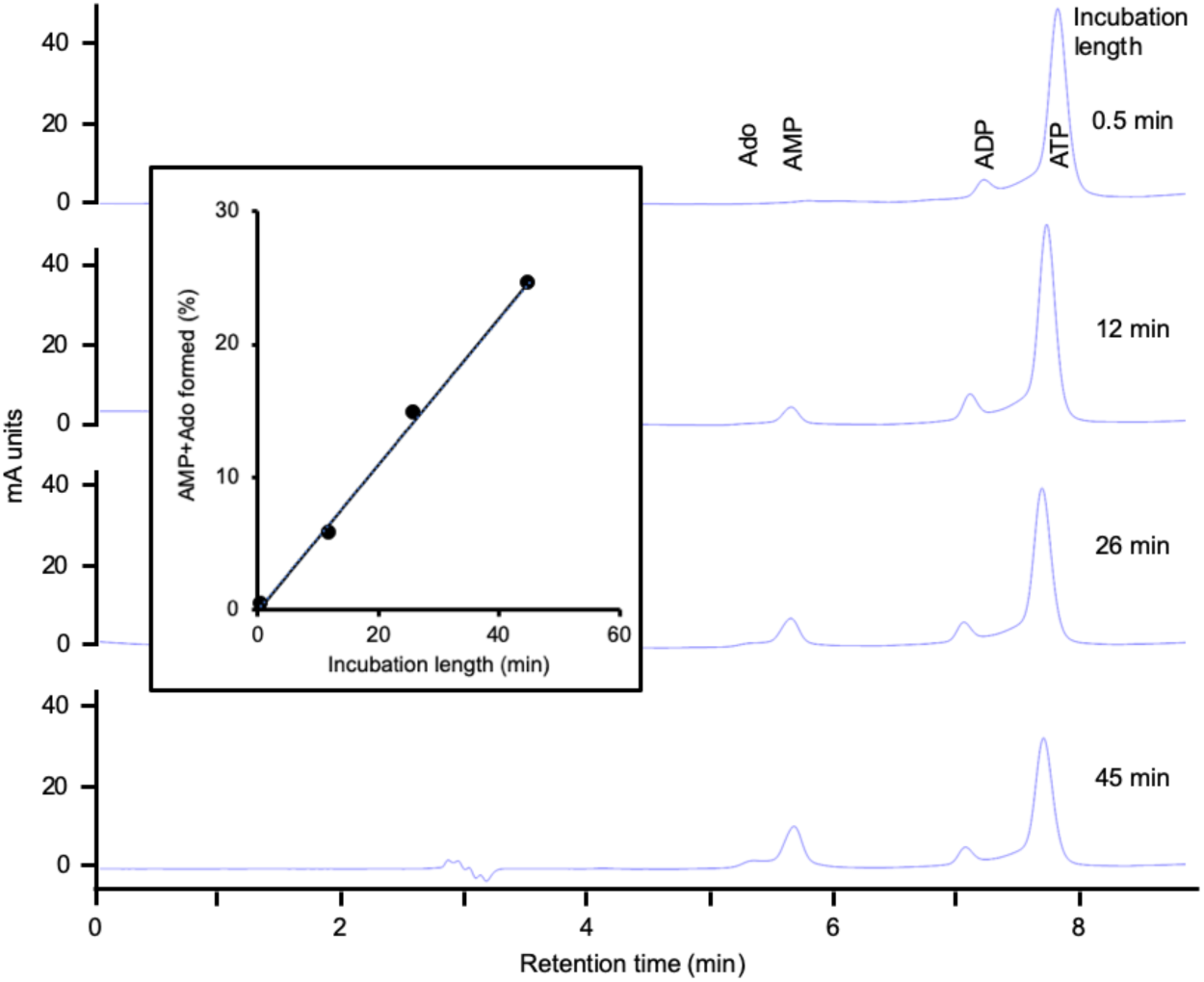
RLNPP/PDE hydrolysis of ATP. A reaction mixture of 200 µL containing 50 mM Tris-HCl, pH 9, 100 µM ATP and 1 µL of RLNPP/PDE was incubated at 37°C. At the indicated times, 20 µL were analyzed by HPLC. The insert shows the quantitative results of the reaction mixture in terms of AMP+Ado formed. AMP was a direct product of RLNPP/PDE and Ado was a secondary product due to a contaminating nucleotidase activity

The comparisons, summarized in Table 5, indicate that the best candidate to define a molecular identity for RLNPP/PDE is Enpp3, which is fully compatible with the activities considered. Enpp4 and Enpp5 can be discarded by the absence of significant activity towards ATP and 2’,3’-cGAMP of the human and murine orthologs, and in the case of Enpp5 also by the absence of activity on 4-nitrophenyl-dTMP. Smpdl3a can be discarded by the acid pH optima of the activities on ATP and 4-nitrophenyl-dTMP. However, with the only data shown in Table 5, it cannot be discarded that a very minor part of the activity of RLNPP/PDE on 4-nitrophenyl-dTMP and/or on 2’,3’-cGAMP at pH 9 (as in the assays of the amino acid effect on the inactivation by EDTA, and of 2’,3’-cGAMP hydrolase) could be due to Enpp4 or Smpdl3a, respectively. Nevertheless, the activity of Enpp4-like enzymes on 4-nitrophenyl-dTMP is not documented other than by an incidental mention (Albright et al. 2012) and that on ATP is considered negligible (Albright et al. 2014). On the other hand, the presence of significant Smpdl3a activity in RLNPP/PDE is not supported by the very low activity towards 4-nitrophenyl-phosphorylcholine as compared to that on 4-nitropheyl-dTMP (Table 3), as both substrates are hydrolyzed at the same rate by human SMPDL3A (Traini et al. 2014). Therefore, most part, if not all of the phosphodiesterase activity of RLNPP/PDE on 4-nitrophenyl-dTMP (whose inactivation by EDTA was accelerated by amino acids) as well as the 2’,3’-cGAMP hydrolase activity, are due to rat Enpp3.

**Table 5.** Comparison of RLNPP/PDE activities with those of relevant enzymes detected in RLNPP/PDE by LC-MS/MS.

| Enzyme | Nucleotide pyrophosphatase on ATP | Phosphodiesterase on 4-nitrophenyl-dTMP | Phosphodiesterase on 2',3'-cGAMP |
| --- | --- | --- | --- |
| RLNPP/PDE | Active at pH 7.5–8.0 (Ribeiro et al. 2000) and at pH 9 (Fig. 7) | Active with alkaline pH optimum (Fig. S1C) | Active at pH 9 (Fig. 6) |
| Enpp3 | Rat enzyme is active at pH 9.5 (Dohler et al. 2018) | Rat enzyme is active with alkaline pH optimum (Deissler et al. 1995) | Human and murine orthologs are active at pH 4–9 (Mardjuki et al. 2024) |
| Enpp4 | Human ortholog shows very minor activity (Albright et al. 2014) | Human ortholog is active at unknown pH (Albright et al. 2012) | Murine ortholog shows negligible activity (Mardjuki et al. 2024)<sup>1</sup> |
| Enpp5 | Human and murine orthologs show negligible activity (Gorelik et al. 2017) | Rat enzyme shows negligible activity (Ohe et al. 2003) | Murine ortholog shows negligible activity (Mardjuki et al. 2024)<sup>1</sup> |
| Smpdl3a | Human and murine orthologs are active with acid pH optimum (Gorelik et al. 2016; Traini et al. 2014) | Human ortholog is active with acid pH optimum (Traini et al. 2014) | Murine ortholog is active at pH 4–10 (Hou et al. 2023) |

<sup>1</sup> Enpp1 and Enpp3 account for all the 2',3'-cGAMP hydrolase activity in mouse cells

### Essentiality of the α-amino-carboxyl structure for the effects of amino acids on the acceleration of RLNPP/PDE inactivation by EDTA

Comparisons of selected amino acids highlight simple structural features of the R group that diminish the acceleration factor or AF value (Table 1 and Fig. 3), including the following cases (AF values from Table 1 are given in parenthesis). The addition of a methylene group produced in most cases a ≈1.5-fold decrease of AF value, as indicated by comparison of glycine (6.6) with alanine (3.7), aspartate (5.3) with glutamate (3.4), and asparagine (3.7) with glutamine (2.7). An exception to this rule comes from the comparison of valine (2.0) with leucine (2.1) or isoleucine (1.8), perhaps due to the ramified carbon chain of the R group. The amidation of a carboxyl group produced a ≈1.3-fold decrease of AF value, as indicated by comparison of aspartate (5.3) with asparagine (3.7), and glutamate (3.4) with glutamine (2.7). The addition of an aromatic ring produced also a ≈1.3-fold decrease of AF value, as indicated by comparison of alanine (3.7) with phenylalanine (3.1), and serine (4.4) with tyrosine (3.2). The removal of a hydroxyl group produced a 1.2–1.7-fold decrease, as indicated by comparison of serine (4.4) with alanine (3.7), and hydroxyproline (2.7) with proline (1.6).

Comparisons of amino acids with analogs in which the α-amino-carboxyl functional group is modified, highlight the essential character of this structure, including the following cases (AF values from Table 1, for L-amino acids, or Table 2, for analogs, are in parenthesis). The removal of the α-amino group produced a strong, 3–5-fold decrease of AF as indicated by comparison of glycine (6.6), aspartate (5.3), alanine (3.7) and glutamate (3.4) with their respective deaminated compounds acetate (1.3), succinate (1.1), propionate (1.1) and glutarate (1.1). The substitution of the α-amine by an α-oxo group produced a strong, 2–4-fold decrease of AF as indicated by comparison of aspartate (5.3), alanine (3.7) and glutamate (3.4) with their respective α-oxo derivatives oxaloacetate (1.2), pyruvate (1.6) and α-ketoglutarate (1.4). The substitution of the α-amine by an α-hydroxyl group, produced a strong, 3–5-fold decrease of AF as indicated by comparison of aspartate (5.3) and alanine (3.7) with their α-hydroxy derivatives L-malate (1.1) and L-lactate (1.2). The removal of the α-carboxyl group produced a strong, 2.5–5.5-fold decrease of AF as indicated by comparison of glycine (6.6) and alanine (3.7) with their decarboxylated derivatives aminomethane (1.2) and aminoethane (1.2). Increasing the covalent distance between the carboxyl and amine groups produced a strong, 4–7-fold decrease of AF as indicated by comparison of glycine (6.6) to glycylglycine (1.5), glycylglycylglycine (1.2), β-alanine (0.9), γ-aminobutyrate (1.0) and ε-aminohexanoate (1.1).

In summary, the α-amino-carboxyl structure typical of the common amino acids is necessary to accelerate RLNPP/PDE inactivation by EDTA. On the other hand, the presence of an imino instead of the amino group produced little effect on AF as indicated by the comparison of proline (1.6) or hydroxyproline (2.7) with for instance the aliphatic amino acids (1.8–2.1). It seems likely that the D configuration of amino acids is similarly efficient as the L one, since the AF values obtained with D-alanine (2.3) and D-aspartate (5.3) were like those observed with L-alanine (3.7) and L-aspartate (5.3).

### With a few exceptions, amino acid size correlates negatively, and stability constant of amino acid-Zn^2+^ complex correlates positively with the acceleration of RLNPP/PDE inactivation by EDTA

Comparisons among amino acids indicated that some features that represent augmentation of the R group (addition of methylene or aromatic ring, carboxyl amidation) diminished the AF value. Therefore, correlation analyses were performed between the increase of k_i(ap)_ and two size-related parameters (crude data listed in Table S2). To analyze the correlation with respect to molecular weight, all the amino acids studied in Fig. 3 were considered, which yielded a very weak, non-significant negative correlation (Fig. 8, left panel). To analyze the correlation with respect to partial molar volume, glutamine and tyrosine could not be considered due to lack of data. In this case the negative correlation was somewhat better but still weak and non-significant (Fig. 8, central panel). Nevertheless, in both correlation analyses, the point corresponding to tryptophan was clearly separated from the rest of amino acids. Therefore, the correlations were reanalyzed omitting tryptophan data. The resulting negative correlations were stronger and significant, particularly with partial molar volume (Fig. 8, left and central panels; see data shown in red color).

**Fig. 8.**
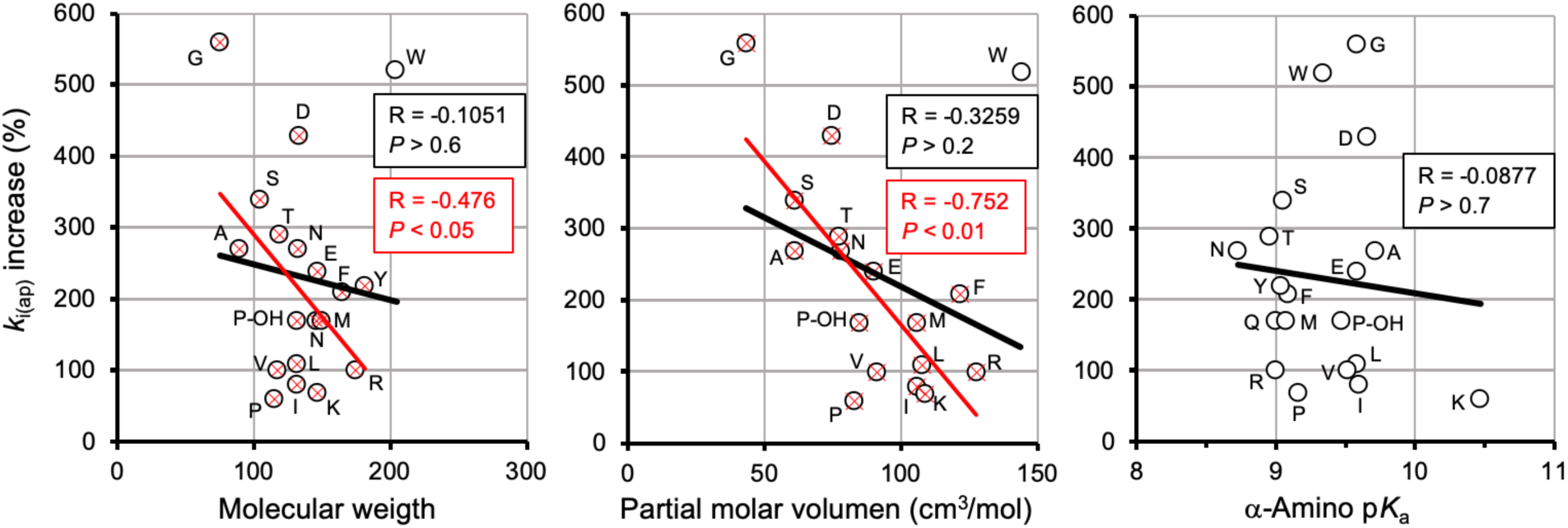
Correlation analyses of k_i(ap)_ increase with amino acid parameters: molecular weight, partial molar volume and α-amino pK_a_. Molecular weights were taken from PubChem database (Kim et al. 2023); partial molar volumes and pK_a_ values from ((Haynes et al. 2017). Left and center panels show two datasets: (i) circles and black regression line (all the amino acids for which data were available) and (ii) red crosses (overlapped with circles) and regression line (same amino acids except one clear outlier which corresponds to tryptophan). The right panel contains a single dataset (all the amino acids for which data were available). The resulting Pearson correlation coefficients and *P* values are indicated in the panels. Amino acids are identified by the one-letter code. P-OH, hydroxyproline.

The study of the amino acid effects on RLNPP/PDE inactivation by EDTA was carried out at pH 9.0, which is near the typical value of the α-amino pK_a_. It was reasoned that this could lead to different ionization degrees of this group, which is essential for the effect of amino acids (see above). Therefore, the possible correlation between the increase of k_i(ap)_ and the α-amino pK_a_ value was also analyzed (Fig. 8, right panel; crude data listed in Table S2), but the results indicated the absence of any significant relationship even when some outliers were omitted (not shown).

It seemed likely that the pro-inactivating effect of amino acids would involve somehow an interaction with Zn^2+^ ions. Therefore, correlation analyses were also run between the increase of k_i(ap)_ and the stability constants of free amino acid-Zn^2+^ complexes. In the NIST database (Smith et al. 2004), stability constants are recorded at different temperatures and ionic strength, but data for all amino acids under the same conditions are not available. Two different datasets were composed (see Table S2) by choosing constants either at 37°C and ionic strength (µ) 0.15 (no data for hydroxyproline and tyrosine) or at 25°C and µ 0.10 (no data for glutamine, glutamate and isoleucine). The correlations obtained with both datasets are shown in Fig. 9. Both cases gave similar results. A weak, non-significant positive correlation was observed when all the datapoints were considered (Fig. 9, both panels; see data colored black). However, after removing two or three very clear outliers in each case (corresponding to aspartate, proline and hydroxyproline), strong and significant positive correlations were apparent (Fig. 9, both panels; see data shown in red color).

**Fig. 9.**
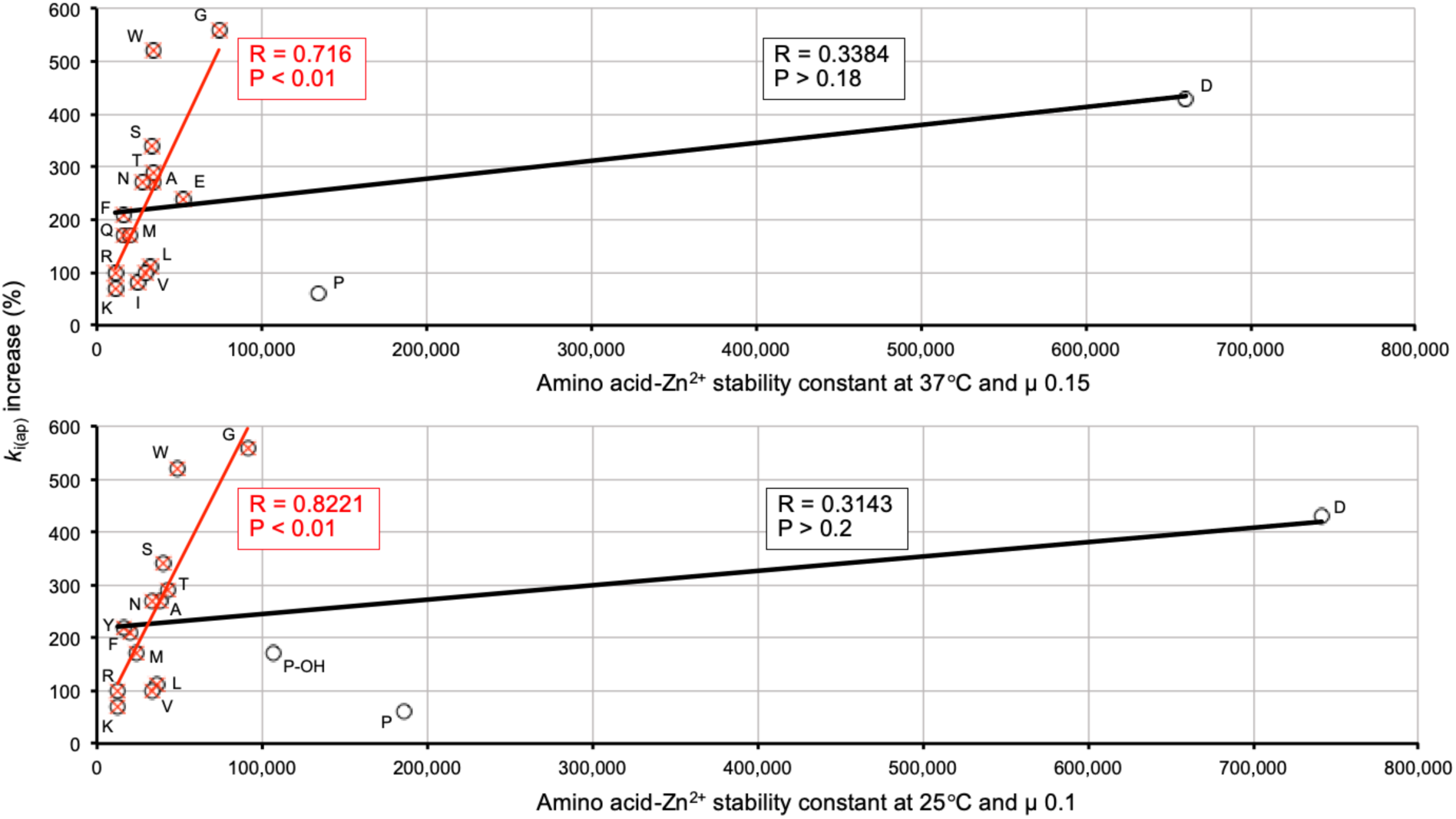
Correlation analyses of k_i(ap)_ increase with amino acid-Zn^2+^ stability constants. The stability constants of amino acid-Zn^2+^complexes, at either 37°C and µ 0.15 (upper panel) or at 25°C and µ 0.10 (lower panel), were taken from (Smith et al. 2004). Each panel shows two datasets: (i) circles and black regression line (all the amino acids for which stability constants were available) and (ii) red crosses (overlapped with circles) and regression line (same amino acids except two or three clear outliers). The numerical data used in these plots are shown in Table S1. The resulting Pearson correlation coefficients and *P* values are indicated in the panels. Amino acids are identified by the one-letter code. P-OH, hydroxyproline.

In summary, the correlations shown in Fig. 8 and Fig. 9 establish a combined relationship, whether causal or not, of the acceleration effect evoked by amino acids with their size and with their affinity for Zn^2+^. In general, the smaller the amino acid and the higher its affinity for Zn^2+^, the stronger is the acceleration produced of the time-dependent inactivation of RLNPP/PDE by EDTA. However, a few meaningful exceptions are worth of note. In Fig. 8 (left and central panels), tryptophan, which is the largest amino acid displays one of the stronger effects, only slightly below glycine which is the smallest amino acid. In Fig. 8 (both panels), aspartate, which is by far the amino acid with the highest affinity for Zn^2+^, accelerates enzyme inactivation by EDTA somewhat less than glycine and tryptophan. Also exceptional were proline and hydroxyproline, which produced lower k_i(ap)_ increases than their relatively high affinities for Zn^2+^ would predict. All these exceptions indicate that the effect of the amino acids cannot be rationalized on a single parameter.

### Mechanistic interpretation of the effect of free amino acids as accelerators of the time-dependent inactivation of RLNPP/PDE by EDTA: evidence favors the occurrence of a conformational change evoked by amino acids

The acceleration elicited by free amino acids that themselves do not inhibit or inactivate RLNPP/PDE (Fig. 3 and Fig. 5) can occur in principle through two different mechanisms. In one case, amino acids would bind the enzyme in one (or more) sterically-delimited sites (possibly including contact with enzyme-bound Zn^2+^ through the α-amino-carboxyl structure) and evoke a conformational change that loosens enzyme-bound Zn^2+^ and/or makes it more accessible to EDTA. Another mechanism that could take place would be that, rather than binding to RLNPP/PDE, amino acids could bind in solution to EDTA-Zn^2+^ complexes formed from Zn^2+^ slowly released from the enzyme. The formation of the ternary EDTA-Zn^2+^-amino acid complexes would stabilize and possibly accelerate Zn^2+^ chelation.

Both mechanisms are not mutually exclusive and can perhaps coexist. However, the experiments shown in Fig. 6 argue strongly against the formation of EDTA-Zn^2+^-amino acid complexes in solution as the only explanation for the pro-inactivating effect of amino acids. In that experiment, the effect of EDTA alone was independent on the time elapsed from membrane preparation to assay, whereas the acceleration evoked by glycine increased with time. There is no reason why the formation of EDTA-Zn^2+^-amino acid complexes in solution should be affected by the time elapsed after membrane preparation. Therefore, we conclude that there is a direct interaction of free amino acids with RLNPP/PDE.

### About the possible physiological relevance of an effect of amino acids on the conformation of Enpp3

Mammalian Enpp3 enzymes are multidomain integral membrane proteins. Most of their structure, after small cytosolic (CD) and transmembrane (TM) domains, faces the extracellular medium containing two N-terminal tandem somatomedin B-like domains (SMB), a phosphodiesterase domain (PDE) and a C-terminal nuclease-like domain (NUC) (Borza et al. 2021). Most of our experiments were performed using the RLNPP/PDE solubilized by limited trypsinization under native conditions, which likely corresponds to the Enpp3 protein devoid of the CD and TM domains. Extrapolation of the results to enzyme in its physiological membrane location is supported by experiments demonstrating that the amino acid-dependent acceleration of the time-dependent inactivation of RLNPP/PDE by EDTA takes place also in crude membranes. The effect was observed with freshly obtained membranes, but its intensity increased when tests were delayed a few days after membrane preparation (Fig. 6).

In its natural location facing the extracellular medium, Enpp3 is exposed not to a single amino acid, but to mixtures typical of serum which fluctuate depending on nutritional status. Tests run with amino acid mixtures at different concentrations (0.25–4-fold those prevailing in portal blood of fasted rats; Fig. 5) showed that the amino acid effect on Enpp3 in vivo may depend on the nutritional status of the animal.

### Roles of Enpp1 and Enpp3 orthologs dependent or independent on enzyme activity and on protein contacts: potential relevance of a conformational change evoked by amino acids

Although the results of this work are primarily related to Enpp3, the functions of both Enpp1 and Enpp3 ortholog proteins are discussed here, as both enzyme kinds display a marked similitude in their structures and catalytic activities (Borza et al. 2021; Gorelik et al. 2018; Stefan et al. 2005). For this reason, after attributing to Enpp3 the amino acid-dependent acceleration of inactivation by EDTA, one wonders whether the same response could be observed with Enpp1.

Despite their similitude, it must be remarked that Enpp1 and Enpp3 differ in their tissular expression profiles, remarkably with a different subcellular distribution in rat hepatocytes, where Enpp1 is located in the basolateral and Enpp3 in the apical surface (Scott et al. 1997).

Enpp1 orthologs are the most studied members of the ENPP family, with many roles in health and disease. This includes both functions that are dependent or independent on enzyme activity. (i) A well-known example of the former is the regulation of calcification in bone and other tissues susceptible to mineralization. The hydrolysis of extracellular ATP by Enpp1 orthologs generates pyrophosphate (PPi), which is an inhibitor of hydroxyapatite deposition and prevents overmineralization. Deficiency or inactivating mutations of ENPP1 are clearly related to the occurrence of several calcification disorders (Borza et al. 2021; Ferreira et al. 2023; Orriss et al. 2016; Roberts et al. 2019; Terkeltaub 2006; Terkeltaub 2001). (ii) Also dependent on the activity of Enpp1 enzymes is their role in purinergic signaling due to the hydrolysis and generation of agonists (Ruiz-Fernández de Córdoba et al. 2023; Stefan et al. 2006). (iii) Another activity-dependent function of ENPP1 is its immunoregulatory role through the cGAS (2’,3’-cGAMP synthase)-STING pathway (Decout et al. 2021), where it intervenes by hydrolyzing 2’,3’-cGAMP (Carozza et al. 2022; Li et al. 2014). This has stimulated the search of ENPP1 inhibitors with potential application in cancer immunotherapy (Carozza et al. 2020; Cogan and Bakhoum 2020; Onyedibe et al. 2019; Rauf et al. 2023; Ruiz-Fernández de Córdoba et al. 2023). (iv) Finally, an important function attributed to ENPP1 is the inhibition of the insulin receptor tyrosine kinase by ENPP1, which has a role in the insulin resistance of non-insulin-dependent diabetes mellitus (Arianti et al. 2021; Goldfine et al. 1998; Goldfine et al. 2008; Goldfine et al. 1999; Maddux et al. 1995; Roberts et al. 2019; Teno et al. 1999). In this case there is contradictory evidence, still unresolved, for the dependence of the inhibition of the insulin receptor on the enzyme activity of ENPP1 (Chin et al. 2009; Grupe et al. 1995; Stefan et al. 1996), but anyhow such inhibition has been repeatedly related to a direct contact between both membrane proteins (Di Paola et al. ; Dimatteo et al. 2013; Kulesza et al. 2023; Maddux et al. 1995; Tassone et al. 2018).

Enpp3 homologues have also been attributed with multiple roles. This covers functions that are clearly dependent on enzyme activity, as well as some that have not yet been characterized in this regard. (i) Enpp3 is highly expressed in basophils and mast cells and, through its ATP hydrolytic activity, it has been shown to regulate negatively the ATP-dependent chronic allergic responses (Bühring et al. 2001; Tsai et al. 2015). (ii) High Enpp3 expression has also been observed in epithelial cells of the small intestine, and its ATP hydrolytic activity contributes to the maintenance of the interferon-producing plasmocytoid dendritic cells, which are very sensitive to ATP-induced cell death (Furuta et al. 2017). (iii) Another activity-dependent function of ENPP3 is immunoregulation through the cGAS-STING pathway, mediated by the hydrolysis of 2’,3’-cGAMP (Mardjuki et al. 2024), an activity that has been confirmed in this work (Fig. 7). (iv) Evidence links Enpp3 orthologs to the female reproductive system, as it is highly expressed in the uterine epithelium. ENPP3 undergoes cyclic changes in the human endometrium, where it affects embryo adhesion and invasion by modifying implantation-factors expression (Boggavarapu et al. 2016; Chen et al. 2018). In rats, Enpp3 expression has also been related to the morphological changes and inflammatory response during ovulation and luteinization (Li et al. 2017). (v) Enpp3 expression has also been shown to be involved in the invasive properties of tumor cells of glioma (Deissler et al. 1999) and colon carcinoma (Yano et al. 2003), but the mechanism for these effects is unknown. (vi) Finally, there is circumstantial evidence that Enpp3 may be also related to diabetes, though a direct link to the insulin receptor, as that known to occur in the case of ENPP1, has not been reported (Ghanaat-Pour et al. 2007; Goldsworthy et al. 2013; Liu et al. 2019; Ye et al. 2024).

The effects of free amino acids on RLNPP/PDE affect enzyme activity only in the presence of EDTA, a metal chelator. This is most unlikely to have a reflection under physiological conditions. Concerning the conformational change of Enpp3 evoked by amino acids, one should not expect it affects enzyme activity. So, to put these results in a proper context, one can consider speculatively whether such conformational change may be relevant to any Enpp3 role dependent on protein contacts with(out) intervention of its enzyme activity. On the other hand, if the amino acid effect, here reported with Enpp3, were extended to the structurally and enzymatically similar Enpp1, it would be conceivable that the effect of ENPP1 on the insulin receptor were affected by the change of conformation evoked by free amino acids. The same could apply if the above-mentioned relationship of Enpp3 to diabetes were mediated by contact with the insulin receptor. The potential relevance of these hypothesis is highlighted by the known role of free amino acids on the pathogenesis of insulin resistance (Chen et al. 2022; Gar et al. 2018; Kolic et al. 2023; Krebs et al. 2002; Lee et al. 2018; Patti et al. 1998; Saleem et al. 2019; Seibert et al. 2015; Tremblay et al. 2007).

## Acknowledgement

We are grateful to Felipe Clemente, from the Proteomic Unit (CAI Biological Techniques) of Complutense University of Madrid (Madrid, Spain), for his expert advice.

## Author contributions

JMR and JCC conceived, designed and drafted the manuscript, and carried out formal analyses of data. AF, MJC and JCC supervised experiments performed by other coauthors. AR, GCL, AF, JM, RF, JLG and MJC performed experiments and obtained data. All authors evaluated and reviewed the manuscript, and agreed to the submission.

## Data availability

The data presented in this study are available in the article and supplementary material. Further inquiries can be directed to the corresponding authors.

