## Supplemental Table S2 for "Free Amino Acids Accelerate the Time-Dependent Inactivation of Rat Liver Nucleotide Pyrophosphatase / Phosphodiesterase Enpp3 elicited by EDTA"

**to**

### Free Amino Acids Accelerate the Time-Dependent Inactivation of Rat Liver Nucleotide Pyrophosphatase / Phosphodiesterase Enpp3 elicited by EDTA

**By**

**Ana Romero, Guadalupe Cumplido-Laso, Ascensión Fernández, Javier Moreno, José Canales, Rui Ferreira, Juan López-Gómez, João Meireles Ribeiro, María Jesús Costas and José Carlos Cameselle**

Table S2. Data used for the correlation analyses shown in main manuscript Fig. 8 and Fig. 9.

| Amino acid | $k_{i(ap)}$ increase (%) | Molecular weight | Partial molar volumen ($cm^3$/mol) | α-Amino group p$K_a$ | Amino acid-$Zn^{2+}$ stability constant (25°C, μ 0.1) | Amino acid-$Zn^{2+}$ stability constant (37°C, μ 0.15) |
|---|---|---|---|---|---|---|
| Alanine | 270 | 89.09 | 60.54 | 9.71 | 38018.94 | 34673.69 |
| Arginine | 100 | 174.20 | 127.42 | 9.00 | 12589.25 | 11220.18 |
| Asparagine | 270 | 132.12 | 78.00 | 8.73 | 33113.11 | 28183.83 |
| Aspartic acid | 430 | 133.10 | 74.80 | 9.66 | 741310.24 | 660693.45 |
| Glutamine | 170 | 146.14 | – | 9.00 | – | 16595.87 |
| Glutamic acid | 240 | 147.13 | 89.85 | 9.58 | – | 52480.75 |
| Glycine | 560 | 75.07 | 43.26 | 9.58 | 91201.08 | 74131.02 |
| Isoleucine | 80 | 131.17 | 105.80 | 9.60 | – | 25118.86 |
| Leucine | 110 | 131.17 | 107.77 | 9.58 | 36307.81 | 32359.37 |
| Lysine | 70 | 146.19 | 108.50 | 9.16 | 12882.50 | 11481.54 |
| Methionine | 170 | 149.21 | 105.57 | 9.08 | 23988.33 | 19952.62 |
| Phenylalanine | 210 | 165.19 | 121.50 | 9.09 | 20417.38 | 16595.87 |
| Proline | 60 | 115.13 | 82.76 | 10.47 | 186208.71 | 134896.29 |
| Hydroxyproline | 170 | 131.13 | 84.49 | 9.47 | 107151.93 | – |
| Serine | 340 | 105.09 | 60.62 | 9.05 | 39810.72 | 33884.42 |
| Threonine | 290 | 119.12 | 76.90 | 8.96 | 42657.95 | 34673.69 |
| Tryptophan | 520 | 204.22 | 143.80 | 9.34 | 48977.88 | 34673.69 |
| Tyrosine | 220 | 181.19 | – | 9.04 | 16595.87 | – |
| Valine | 100 | 117.15 | 90.75 | 9.52 | 33113.11 | 29512.09 |
